# Visible Light-Controllable Surgical Hemostatic Adhesives Enabled by Vitamin B_12_-Stabilized Poly(α-Lipoic Acid)

**DOI:** 10.1101/2025.09.09.675035

**Authors:** Yutong Huang, Qikun Yi, Shiying Wang, Caihong Deng, Lingling He, Jiali Ma, Mengran Zhang, Hao Tan, Ping Li, Fei Sun

## Abstract

Water-resistant adhesives are indispensable for biomedical applications, from surgical wound closure to internal hemostasis. While poly(α-lipoic acid)-based adhesives show promise for their strong underwater adhesion, their clinical utility has been limited by uncontrolled polymerization-depolymerization dynamics. Here we present a visible light-controllable adhesive system enabled by a simple mix of α-lipoic acid (αLA) and vitamin B_12_ (i.e., adenosylcobalamin or AdoB_12_)—a photolabile compound—without resorting to any chemical modification. Benign visible light irradiation facilitates the cleavage of the C-Co bond and the subsequent formation of B_12_-thiolate complexes that stabilize αLA polymers against depolymerization. The photoresponsive αLA-AdoB_12_ adhesive system has proven highly effective in a wide range of surgical applications, achieving (1) strong bonding in porcine skin models in vitro, (2) reliable sealing of punctured organs (lung, heart and stomach) ex vivo, and (3) immediate hemostasis in active bleeding models such as topical, esophageal, and intestinal wounds in vivo. This work highlights a simple yet powerful strategy for creating visible light-controllable bioglues well-suited for diverse surgical applications.

## Introduction

Water-resistant surgical adhesives with precisely tunable bonding properties could revolutionize operative techniques (*1-3*). Conventional polymeric tissue adhesives face at least three critical limitations: 1) restricted applicability across diverse tissue types due to wet-susceptibility, 2) inability to modulate adhesion post-application, and 3) reliance on petrochemical-derived, non-biodegradable components that raise biocompatibility concerns (*4-6*). These constraints underscore the need for next-generation bioadhesives that combine broad clinical utility with on-demand water-resistant adhesiveness in physiological environments.

α-Lipoic acid (αLA)-based adhesives, leveraging the dynamic disulfide bonds of 1,2-dithiolane, enable reversible polymerization that confers self-healing, stimuli-responsiveness, and closed-loop recyclability (*7, 8*). However, their application has been hindered by spontaneous depolymerization driven by terminal thiols (-SH) or sulfur radicals (-S∙), which destabilizes the αLA polymers. While several chain-end capping strategies using electrophiles or radical scavengers have been developed (*9-13*), these often require complex synthetic modifications and/or harsh reaction conditions, undermining both biocompatibility and clinical feasibility.

To overcome these limitations, we developed a facile photoresponsive chain-end stabilization strategy that operates under ambient conditions—eliminating the need for high temperatures, UV light, or chemical modifications. By integrating adenosylcobalamin (AdoB_12_)—the physiologically active form of vitamin B_12_ with a green-light-sensitive C–Co bond (λ∼510 nm) —into the αLA system, we created an entirely nutrient-based adhesive with optical control over its water-resistant adhesion. Upon visible light exposure (< 550 nm), AdoB_12_ undergoes the cleavage of its C-Co bond, followed by the S-Co coordination with poly(αLA) terminal thiols, which effectively quenches the closed-loop depolymerization (*14-16*) (Fig. 1A). The resulting αLA-AdoB_12_ system has not only demonstrated robust light-activated tissue bonding capabilities in vitro and ex vivo, but also enabled rapid, light-dependent hemostasis in both topical wounds and endoscopic gastrointestinal bleeding in vivo. By combining the two commercially available nutritional compounds to create optically controlled, water-resistant adhesives, this technology offers a transformative solution applicable across a wide clinical spectrum, spanning from emergency field care to complex surgical settings.

**Fig. 1.**
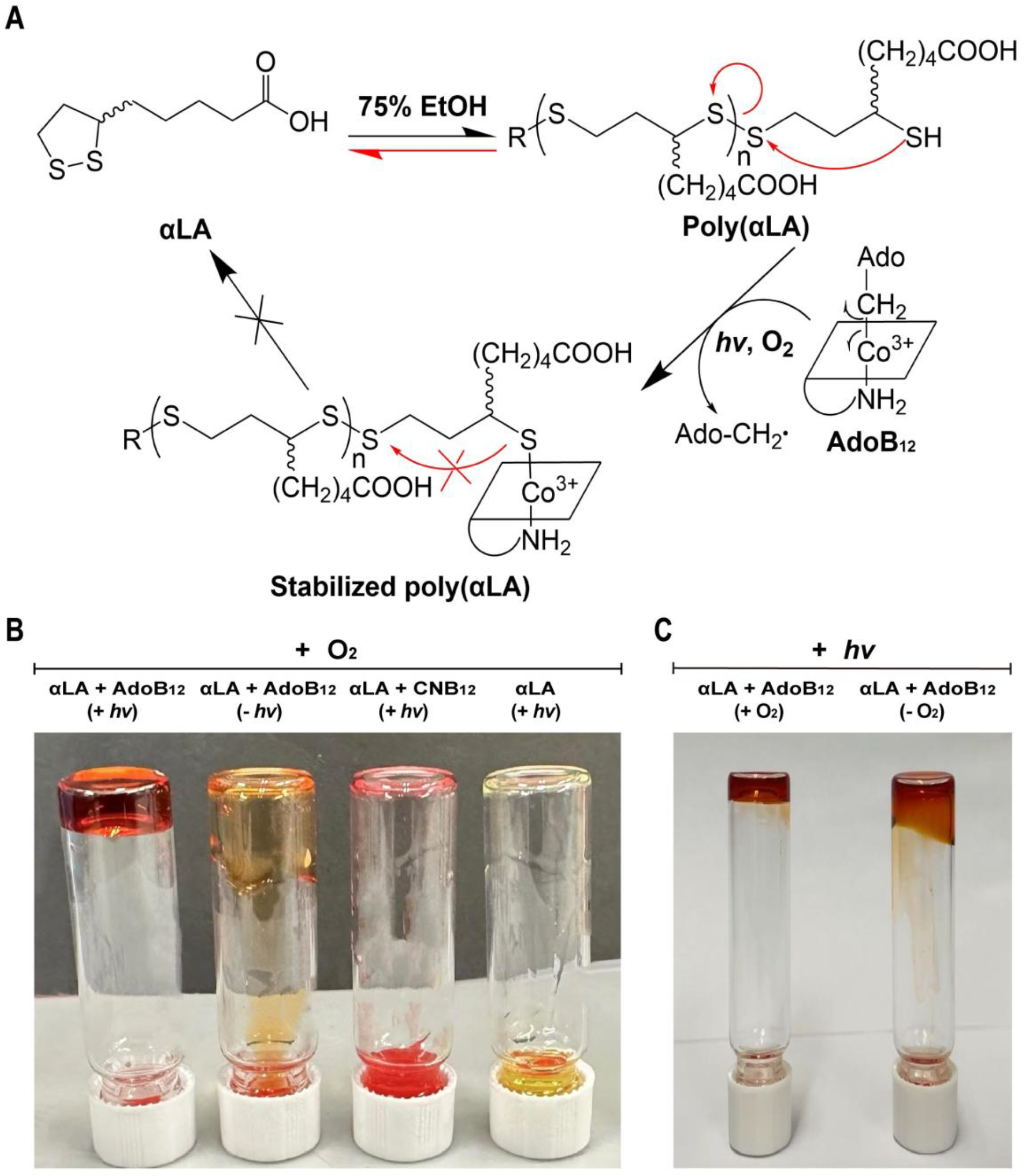
Stabilization of α-lipoic acid polymer by photolysis of adenosylcobalamin (AdoB_12_) under aerobic conditions. (**A**) Scheme of spontaneous αLA polymerization and depolymerization in 75% ethanol and light-induced AdoB_12_-mediated polymer stabilization. White or green light (< 550 nm) induces the cleavage of the Co–C bond and the subsequent formation of a B_12_-thiolate complex, thus halting depolymerization. (**B**) Photographs of αLA solutions (400 mg/mL in 75% ethanol) under ambient conditions. The presence of photolabile AdoB_12_ (10 mg/mL), but not photo-inert CNB_12_, is essential for rapid αLA gelation on exposure to white LED light (20 klux, 5 min). (**C**) Photographs of white light-irradiated αLA–AdoB_12_ solutions (400 mg/mL αLA, 10 mg/mL AdoB_12_ in 75% ethanol) under aerobic and anaerobic conditions. The anaerobic condition was achieved by overnight nitrogen degassing of the solvent.

## Results and Discussion

### Light-Induced Chain-End Stabilization and Rapid Polymerization

High-concentration ethanolic αLA solutions undergo ring-opening polymerization via intermolecular disulfide exchange, forming poly(αLA) (*7, 17, 18*). However, this polymeric state is metastable and spontaneously depolymerizes presumably due to the chain-end thiol radicals, which has thus prompted the development of several chain-end capping and radical scavenging strategies for polymer stabilization.

AdoB_12_, the physiologically dominant form of vitamin B_12_, has emerged as a versatile optogenetic motif, thanks to its green light-sensitive Co-C bond. The AdoB_12_-binding protein CarH has enabled multiple photoresponsive protein materials (*19-23*), as well as chemo-optogenetic tools for controlling complex cellular signalling (*24-26*). Building on this photochemical paradigm, we postulated that cob(III)alamin, a transient intermediate generated from AdoB_12_ photolysis, could scavenge the terminal sulfur radical via Co-S coordination, thereby stabilizing the αLA polymer. To test this, we prepared ethanolic solutions containing αLA (400 mg/mL in 75% ethanol) and AdoB_12_ (10 mg/mL in 75% ethanol). In the dark, the αLA-AdoB_12_ solution remained liquid for >6 hours. Upon white-light exposure (20 klux), however, the αLA-AdoB_12_ solution rapidly increased in viscosity and formed a gel under ambient conditions within 5 minutes (Fig. 1B). By contrast, the mixture of αLA and photo-inert CNB_12_ remained liquid after similar light irradiation, and so did αLA alone (Fig. 1B). Notably, when αLA was dissolved in 75% ethanol that was thoroughly degassed with N_2_, white light irradiation failed to induce its gelation, corroborating the well-established role of O_2_ in the formation of stable Co-S complexes after photolysis (Fig. 1C and movie S1) (*27*). Together, these results demonstrated the photolysis of AdoB_12_ as a feasible strategy to stabilize αLA polymers under ambient conditions.

### Optically Controllable Mechanical Properties of αLA-AdoB_12_ Polymers

Rheological measurements confirmed that light irradiation significantly impacts the mechanical properties of the αLA-AdoB_12_ polymers. Both the duration and intensity of light exposure directly enhanced the material’s stiffness and stability. Frequency-sweep measurements demonstrated that storage modulus (G′) increased dramatically with irradiation time. After 12 hours at 20 klux, G′ values reached 27-76 kPa, three orders of magnitude larger than that of samples irradiated for only 5 minutes (0.01-0.07 kPa) (Fig. 2, A and E). This enhanced mechanics was further evidenced in the strain-sweep measurements; the 12-hour irradiated gel withstood deformations beyond 100% strain, while the 5-minute sample yielded at approximately 40% strain (Fig. 2B). Light intensity also played a critical role in modulating the viscoelastic properties. A 15-minute exposure to high-intensity light (20 klux) produced a gel with G′ values of 0.27-1.05 kPa, significantly stiffer than those formed under low-intensity light (0.1 klux, G′ = 0.02-0.12 kPa) (Fig. 2, C and F). Stronger irradiation also improved yield strain, with the 20-klux-irradiated gel resisting deformation up to ∼50% strain, compared to only ∼10% strain for the 0.1-klux-irradiated gel (Fig. 2D). Together these results demonstrated the feasibility of using light to fine-tune the mechanical properties of αLA-AdoB_12_ polymers.

**Fig. 2.**
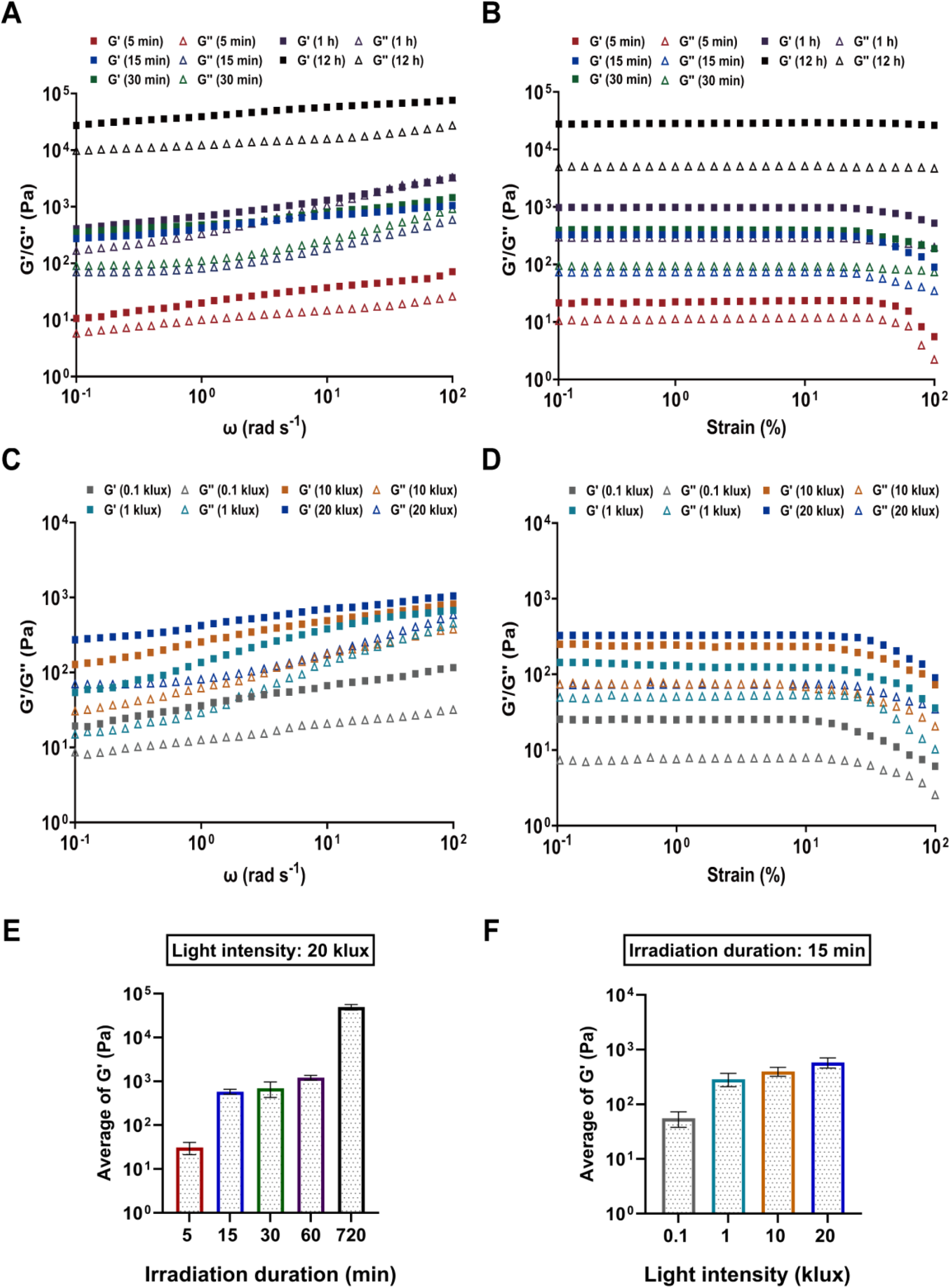
Light-controllable viscoelastic properties of poly(αLA)-AdoB_12_. (**A** and **B**) Frequency-sweep (A) and strain-sweep (B) measurements of gels irradiated at 20 klux for varying durations (5 min to 12 h). (**C** and **D**) Frequency-sweep (C) and strain-sweep (D) measurements of gels irradiated for 15 min at varying intensities (0.1 to 20 klux). (**E** and **F**) Influence of irradiation duration (E) and light intensity (F) on the gel’s stiffness (average G′ over the frequency range of 0.1-100 rad/s). Data are mean ± SD (*n*=3 independent experiments).

Furthermore, the photostabilized poly(αLA)-AdoB_12_ patch exhibited notable self-healing and stretchability, attributable to the dynamic exchange of disulfide bonds within the polymer backbone. Severed films self-healed within 5 minutes at 37°C and retained good stretchability after repair (fig. S1 and movie S2). This unique combination of optically tunable mechanics and intrinsic self-repair capability is critical for applications in dynamic physiological environments—such as pulsatile blood vessels or mobile visceral organs (28, 29)—positioning these materials as promising candidates for demanding clinical applications.

### In Vitro Water-Resistant Skin Adhesion by Photostabilized Poly(αLA)-AdoB_12_

We systematically evaluated the adhesive strength of αLA-AdoB_12_ through lap shear testing using fresh porcine skin substrates. For initial tests, we prepared poly(αLA)-AdoB_12_ patches through visible light-induced gelation. These pre-gelled poly(αLA)-AdoB_12_ revealed only moderate water-resistant adhesiveness, achieving approximately 30 kPa and 55 kPa after 5 min and 12 h of incubation in PBS at 37°C, respectively (fig. S2). This suboptimal adhesion likely stems from the hydrophobic surface of the pre-gelled patches, which compromises interfacial interactions with wet tissues. Substantial improvement was achieved through in situ polymerization, where direct application of the αLA-AdoB_12_ precursor solution followed by 20-klux white light exposure yielded adhesive strengths of ∼80 kPa shortly after 5 min of light irradiation and incubation in PBS at 37°C, further increasing to ∼150 kPa after 12 h, which rivalled commercial products such as CoSeal and Dermabond Advance in tissue adhesion (*9*) (Fig. 3B).

**Fig. 3.**
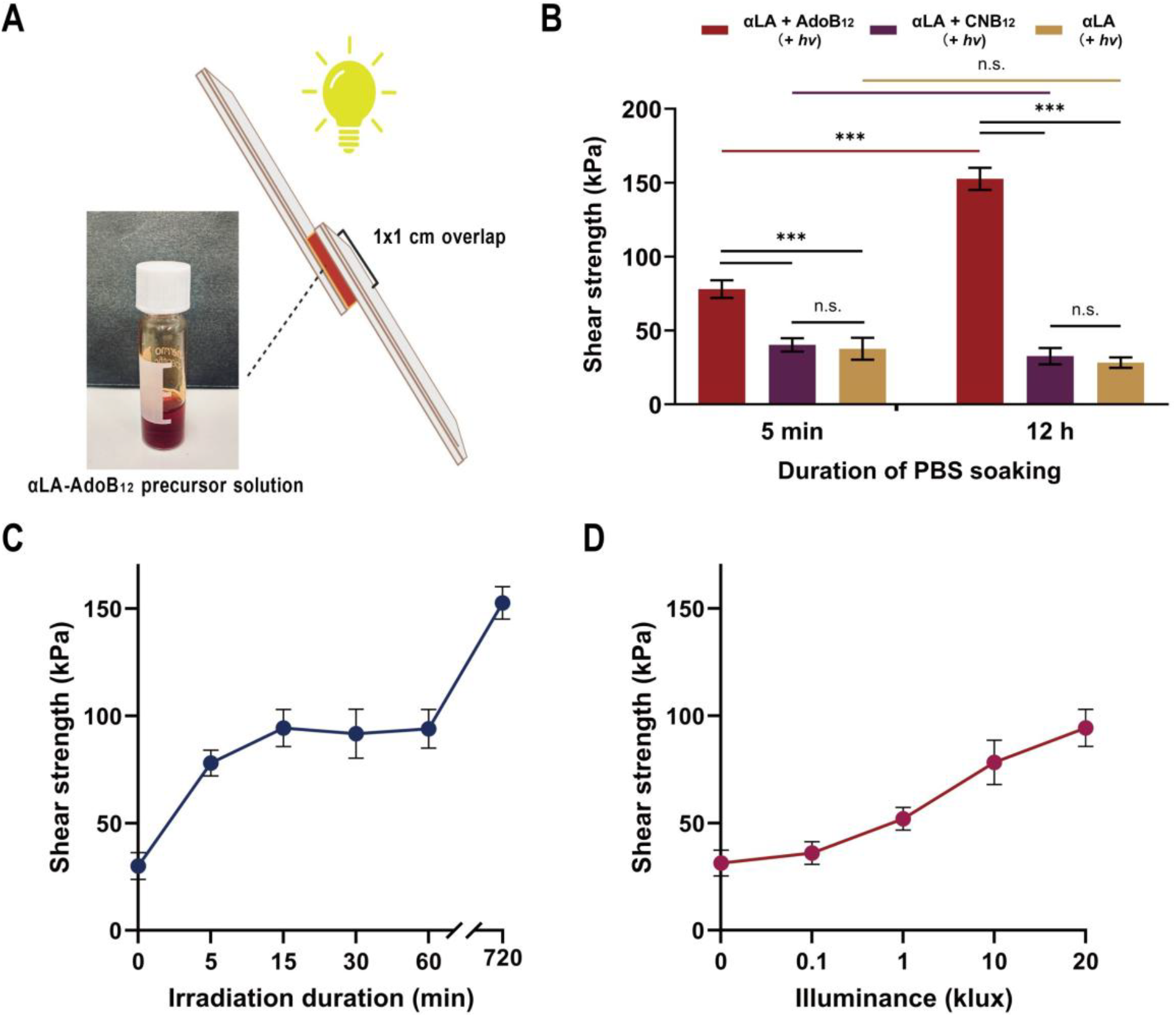
Ex vivo water-resistant adhesive performance of photostabilized poly(αLA)-AdoB_12_. (**A**) Schematic of the lap shear test on wet porcine skin. (**B**) Lap shear strength of various in situ polymerized formulations after 5-min or 12-h PBS soaking at 37 °C. *hv*, white LED light (20 klux). Statistical significance was determined by unpaired two-tailed t-test (*n*.*s*., not significant; ***p ≤ 0.001). (**C** and **D**) Lap shear strength as a function of irradiation duration (constant 20-klux light intensity) (C) and light intensity (constant 15-min irradiation duration) (D). Data are mean ± SD (*n*=3 independent experiments).

The steady increase in adhesive strength, especially within the first 15-min of light irradiation, shows continuous polymerization and network maturation of αLA-AdoB_12_ under light irradiation (Fig. 3C). By contrast, αLA alone or αLA with photo-inert CNB_12_ showed markedly inferior performance, with an adhesive strength of ∼40 kPa after 5 min and even lower after 12 h, which further confirmed the critical role of photolabile AdoB_12_-mediated polymer stabilization in the strong water-resistant adhesion observed. In addition to duration, light intensity also effectively modulated the adhesive strength. At a fixed 15 min of irradiation, increasing the intensity from 0.1 to 20 klux substantially enhanced the lap shear strength, which reached approximately 100 kPa at 20 klux (Fig. 3D). Collectively, these results illustrated light irradiation as a powerful and facile method for controlling the adhesion of αLA-AdoB_12_ to biological tissues.

### Ex Vivo Water-Resistant Organ Sealing by Photostabilized Poly(αLA)-AdoB_12_

We further evaluated the capability of the αLA-AdoB_12_ adhesive system to seal ex vivo porcine organs, including the lung, heart, and stomach. For these tests, 1-cm incisions were made on fresh organs (fig. S3). We initially tested the in situ polymerization strategy; the liquid αLA-AdoB_12_ precursor solution was applied directly onto the incisions and subsequently irradiated with light. Although successful leakage occlusion was achieved (Fig. 4, B and C, and movie S3-S6), the relatively slow solidification process risked the undesirable spread of the precursor solution beyond the target lesion sites. To address this, we adopted the pre-gelling strategy, involving brief light irradiation (20 klux, 5 min) of the precursor solution within a syringe (fig. S4). This treatment generated a viscous, yet still injectable, adhesive. This approach enabled precise application and immediate leakage blockage upon tissue contact (Fig. 4D). The resulting seals demonstrated robust adhesion, withstanding rigorous hydrodynamic challenges including continuous water flushing and simulated tissue movement (fig. S5, and movie S7-S9). The incisions remained fully sealed with no observable leakage. By contrast, αLA alone failed to establish durable closure, exhibiting severe leakage (fig. S6 and movie S10), which further confirmed the critical role of AdoB_12_-mediated photostabilization in achieving effective tissue bonding under dynamic and wet environments. Taken together, these results demonstrated the enormous potential of the αLA-AdoB_12_ system for surgical hemostasis and sealant applications.

**Fig. 4.**
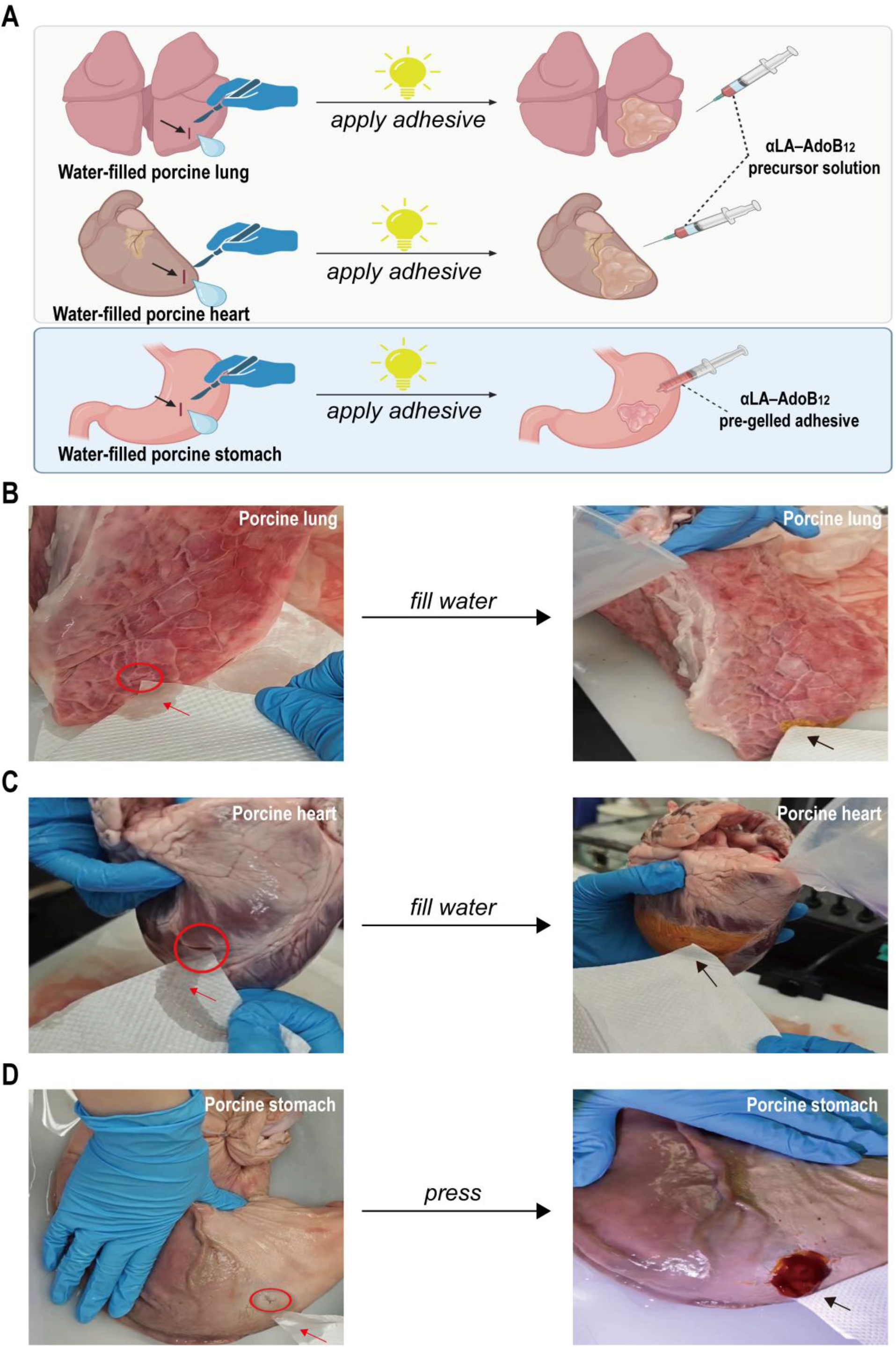
Ex vivo sealing capability of photostabilized poly(αLA)-AdoB_12_. (**A**) Schematic of the sealing tests on water-filled porcine organs (Created with BioGDP.com (*30*)). (**B** and **C**) In situ polymerization of αLA-AdoB_12_ precursor solution for sealing leakage of porcine lung (B) and porcine heart (C). A 1-cm perforation (circled) in water-filled porcine organs leak profusely (red arrow). The perforation is effectively sealed, preventing leakage (black arrow) after continuous water filling. (**D**) Ex vivo sealing demonstration of a porcine stomach by pre-gelled αLA-AdoB_12_ adhesive. A 1-cm perforation (circled) in water-filled porcine stomach leak profusely (red arrow). The perforation is effectively sealed, preventing leakage (black arrow) under mechanical pressure.

### In Vivo Hemostasis in Topical Wounds

Building on successful ex vivo tissue bonding, we evaluated the hemostatic efficacy of the αLA-AdoB_12_ adhesive in a live porcine model with active topical hemorrhage. To ensure rapid sealing, we employed the pre-gelling strategy as aforementioned: the precursor solution was first converted into a viscous, yet injectable, material via brief light irradiation (20 klux, 5 min). This pre-formed gel exhibited rapid, contact-initiated solidification upon blood exposure, forming a conformal seal that achieved immediate hemostasis (Fig. 5, B and C, and movie S11). The formulation possessed balanced physical properties. Its sufficient flowability allowed it to penetrate complex wound geometries without manual packing, while its adequate viscosity resisted washout by blood during application. The resulting adhesive barrier provided immediate and stable hemostasis under physiological stress, thus allowing endogenous clots to mature. It is noteworthy that premature removal of the adhesive with an ethanol-soaked gauze before clot maturation induced rebleeding (movie S12). This result confirmed its primary use as an effective temporary hemostatic sealant for emergency care.

**Fig. 5.**
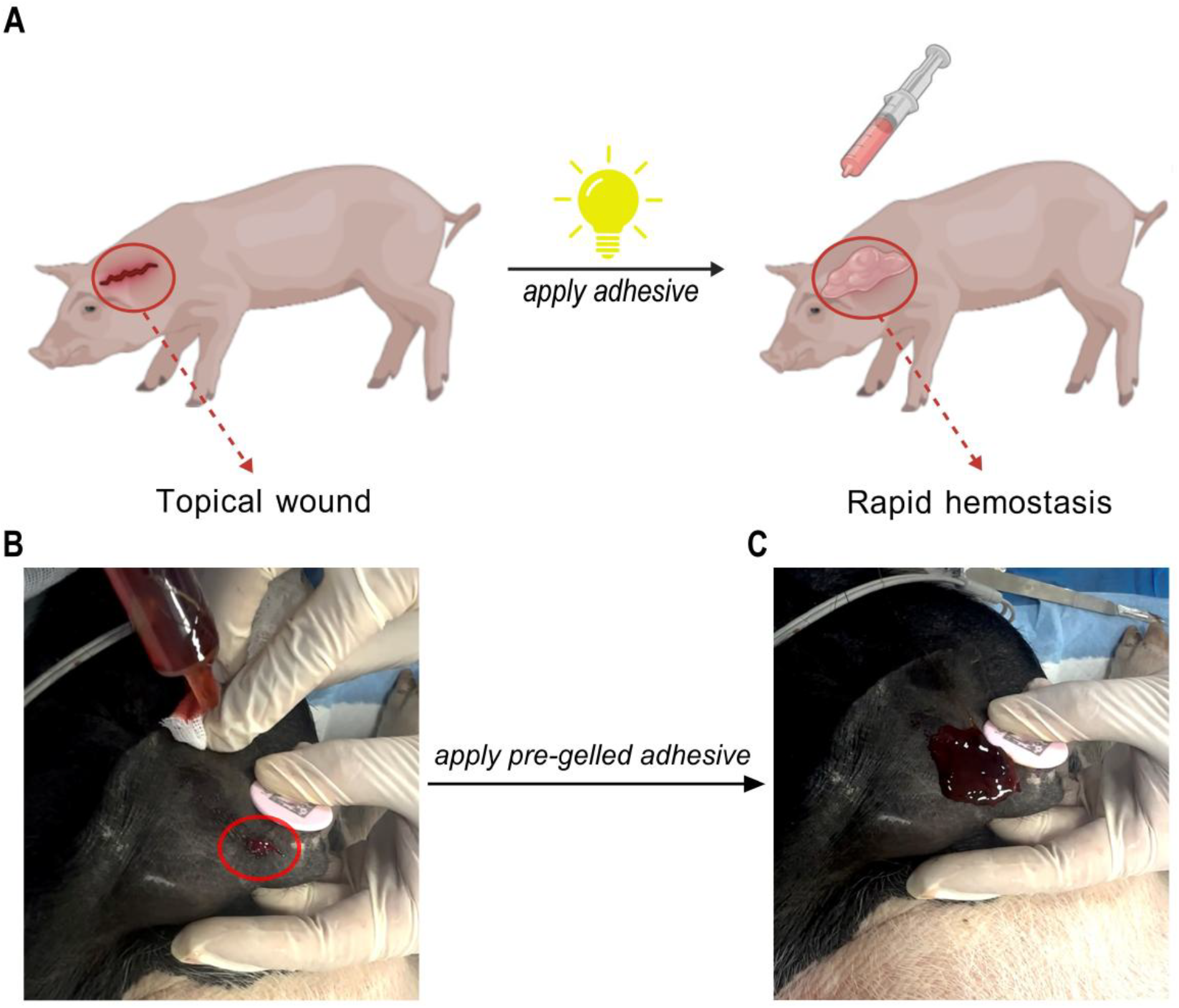
In vivo topical hemostatic performance of the light-activated αLA-AdoB_12_ adhesive in a porcine bleeding model. (**A**) Schematic of the topical wound treatment process: application of the αLA–AdoB_12_ precursor to a bleeding site, followed by light irradiation to achieve hemostasis. (Created with BioGDP.com (*30*)). (**B**) Active hemorrhage (circled) from a surgically created wound prior to treatment. (**C**) Immediate hemostasis achieved upon application of the pre-gelled αLA-AdoB_12_ adhesive, which forms a durable adhesive barrier upon blood contact.

### In Vivo Endoscopic Hemostasis of GI Bleeding

Compared to topical hemorrhage, internal bleeding—such as gastrointestinal (GI) bleeding from peptic ulcerations (the most common cause) or life-threatening variceal hemorrhage—presents far greater therapeutic challenges (*31-33*). Conventional hemostatic agents often fail to maintain strong adhesion under hemodynamic stress, whether on diffuse lesions or large vessel injuries (*34*). This is particularly true for variceal hemorrhage, which remains a critical problem with persistently high rebleeding and mortality rates despite current treatments (*35*). These limitations underscore the urgent need for a generalizable solution capable of managing the full spectrum of internal bleeding.

Our endoscopic validation studies in porcine models demonstrated that photo-stabilized αLA-AdoB_12_ adhesive can effectively address these clinical challenges. Following endoscopic delivery and in situ photopolymerization, the material formed stretchable, blood-resistant seals that withstood physiological blood pressure in both esophageal and gastrointestinal bleeding models (Fig. 6, and movie S13-S14). A brief 20-second light irradiation (20 klux) was sufficient to stabilize the polymer, rapidly halting severe intestinal hemorrhage. This optically controllable adhesive system represents a significant advance over existing endoscopic therapies. Its unique combination of visible light-activated adhesion, immediate flow-resistant hemostasis, and conformal malleability to complex anatomies positions it as a versatile strategy for treating internal bleeding, spanning from minor mucosal leaks to life-threatening arterial hemorrhage.

**Fig. 6.**
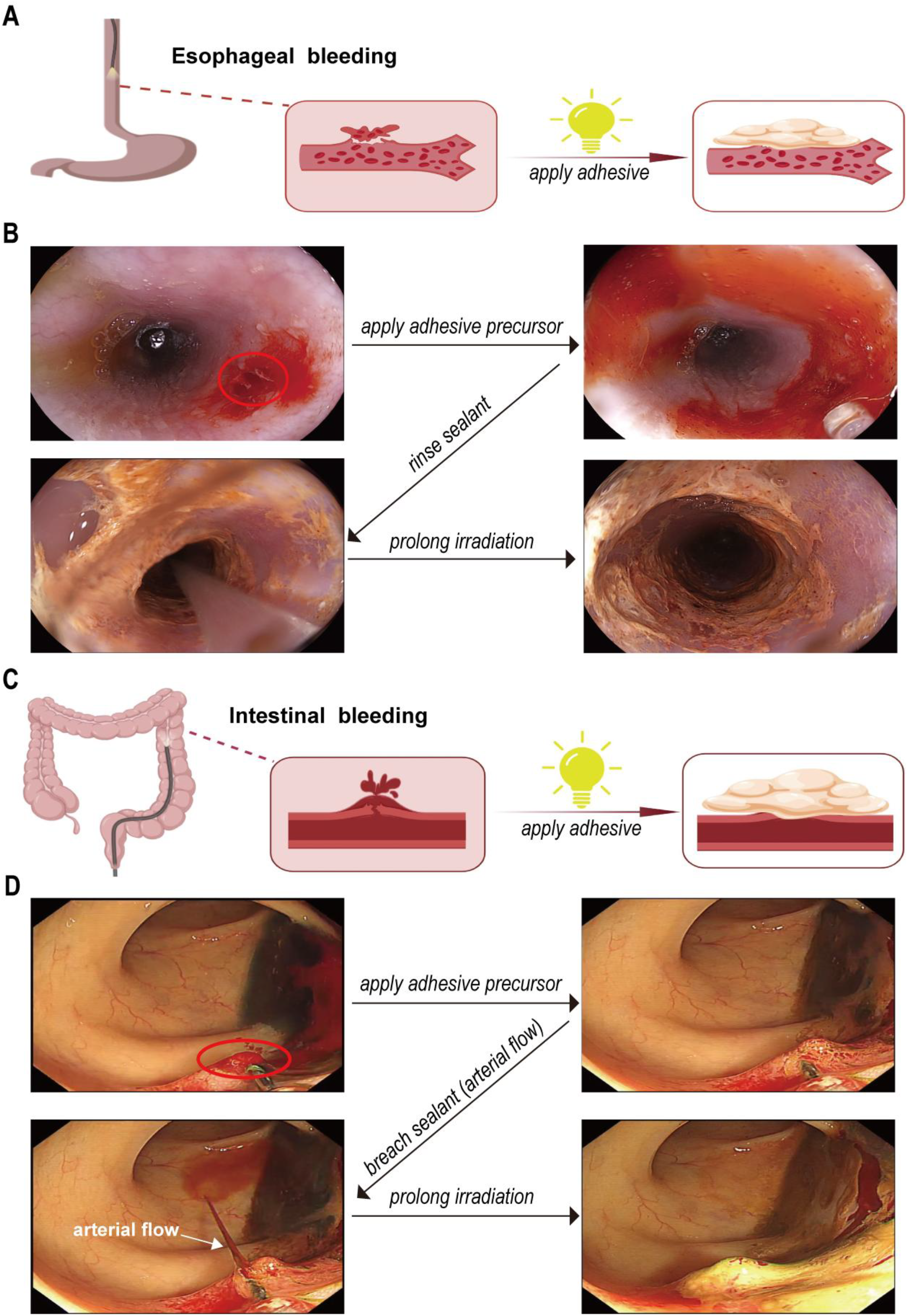
Endoscopic hemostasis of gastrointestinal bleeding using light-activated αLA–AdoB_12_ adhesive in vivo. (**A**) Schematic of treating esophageal mucosal bleeding with the αLA–AdoB_12_ adhesive. (Created with BioGDP.com (*30*)). (**B**) Hemostasis of esophageal mucosal bleeding. Endoscopic delivery of the αLA– AdoB_12_ precursor solution (1 mL in 75% ethanol) to an actively bleeding wound (circled), followed by illumination to form an adhesive sealant. The seal remained adherent under vigorous water flushing; prolonged irradiation further stabilized the seal. (**C**) Schematic of treating arterial intestinal bleeding with the αLA–AdoB_12_ adhesive. (Created with BioGDP.com (*30*)). (**D**) Hemostasis of severe arterial intestinal bleeding. Profuse spurting hemorrhage from a post-polypectomy lesion (circled). Endoscopic delivery of the αLA–AdoB_12_ precursor solution (1 mL in 75% ethanol) and illumination formed an initial seal, which was breached by high-pressure arterial flow (white arrow). Complete and durable hemostasis achieved after reinforcement with a brief 20-sec endoscopic light irradiation (20 klux).

## Conclusion

In summary, we have developed a visible light-controllable adhesive system with broad clinical applicability. Composed of the nutritional compounds αLA and AdoB_12_, the adhesive undergoes rapid stabilization upon visible light irradiation. This process involves photolysis of AdoB_12_ and subsequent coordination of the resulting cobalamin species with terminal thiols of polymerizing αLA, effectively preventing depolymerization. This mechanism confers rapid, light-activated adhesiveness in aqueous and bloody environments, accompanied by high water resistance. The material demonstrates robust tissue bonding and organ sealing ex vivo, as well as immediate and effective hemostasis in vivo—addressing diverse clinical scenarios from topical wounds to active gastrointestinal hemorrhage. Owing to its simple composition and straightforward preparation, this system represents a versatile and practical solution for emergency and surgical applications.

## Author contributions

F.S. conceived the idea and designed the project. Y.H. and Q.Y. performed all polymerization and in vitro analyses. Y.H. and S.W. performed ex vivo experiments. P.L., C.D., L.H., J.M., M.Z., Y.H., and H.T. performed in vivo studies. Y.H., Q.Y., and F.S. wrote the manuscript, and all authors reviewed the manuscript.

## Data and materials availability

All data are available in the main text or the supplementary materials.

## Supplementary Materials

### Materials and Methods

#### Materials

*DL*-α-Lipoic acid (αLA, ≥99%) was purchased from Aladdin (Cat# T106640). Adenosylcobalamin (AdoB_12_, ≥98%) was obtained from Mreda (Cat# M075376). Cyanocobalamin (CNB_12_, ≥98%) was sourced from Sichuan Vicky Biotechnology (Cat# 0001443).

A white LED light source (Philips, 6.5 W, 3000 K, 650 lm) was used for photoirradiation experiments. A photometer (Pro’sKit, MT-4617LED, 0.02–200 klux range, silicon photodiode sensor) was used to measure the light intensity. The following endoscopic instruments were used for the in vivo hemostatic tests: biopsy forceps (RUITIAN, QFB-A-23-1600; diameter: 2.3 mm, length: 1600 mm), a semicircular snare (Micro-Tech, MTN-PFS-A-15/23; diameter: 2.3 mm, length: 2300 mm), and an endoscopic injection needle (Beijing ZKSK Technology, SN18-07/230; 22-gauge, length: 1800 mm).

#### Methods

##### Rheological Measurements

The αLA-AdoB_12_ precursor solutions were cast onto a Parafilm™ M surface in the dark. The samples were then irradiated with white LED light at the specified intensity (0.1, 1, 10, or 20 klux) and duration (5 min, 15 min, 30 min, 1 h, or 12 h) at room temperature. The viscoelastic properties of the resulting materials were measured using a TA Instruments ARES-RFS rheometer equipped with 25 mm and 8 mm parallel steel plates at 23 °C. A gap of 0.5 ± 0.1 mm was maintained between the plates. Frequency-sweep (0.1–100 rad/s, 5% strain) and strain-sweep tests (0.1–100% strain, 1 rad/s) were performed.

##### Lap Shear Tensile Testing on Wet Porcine Skins

1. **Pre-gelled αLA-AdoB**_**12**_ **patch adhesives**. The αLA-AdoB_12_ precursor solution was cast onto a Parafilm™ M surface in the dark. The sample was then irradiated with white LED light at the specified intensity and duration (typically 20 klux for 12 hours) at room temperature. The resulting patch was applied to the 1 × 1 cm overlap area of two porcine skin strips and hand-pressed for 5 sec. The assembled lap joint was incubated in 1× PBS at 37 °C for either 5 min or 12 h. Subsequently, the sample was removed, gently dried with Kimwipes, and subjected to lap shear testing on a TA Instruments ARES-RFS rheometer at a constant displacement rate of 1 mm/min until failure (N = 3). The lap shear strength was calculated by dividing the maximum load at failure by the bonded area.
2. **In situ polymerized adhesives**. A 30 μL aliquot of the αLA–AdoB_12_ precursor solution was evenly applied to the 1 × 1 cm overlap area on wet porcine skin strips. The joint was hand-pressed for 5 sec and immediately immersed in 1× PBS at 37 °C, while being irradiated with white LED light at the specified intensity and duration (typically 20 klux for 12 hours). The sample was gently dried with Kimwipes and then tested on the rheometer under the same conditions (1 mm/min displacement rate, N = 3). The lap shear strength was calculated as described above. Control groups using αLA alone or the photo-inert αLA–CNB_12_ complex were prepared and tested following the same procedures.

##### Ex Vivo Organ Sealing Tests

Fresh porcine lung, heart and stomach were obtained from a local supplier (Freshippo). Organs were thoroughly rinsed with water, and a 1-cm incision was made in each using a scalpel.

1. **Lung and heart**. The αLA–AdoB_12_ precursor solution was applied to the incision site on water-filled organs under white LED light irradiation (20 klux). After 5 min, tissue paper was placed beneath the incision. Additional water was introduced to fully distend the organs, and leakage was assessed. The incision site was flushed with excess water, and organs were pressed manually to evaluate adhesive integrity.
2. **Stomach**. The pre-gelled αLA–AdoB_12_ adhesive was applied to the incision site on a water-filled stomach under white LED light irradiation (20 klux). After 5 min, tissue paper was placed beneath the incision as an indicator of water leakage. The sealed area was subjected to firm manual pressure, and leakage was assessed. The incision site was flushed with excess water, and the stomach was examined manually to evaluate adhesive integrity under hydrodynamic and mechanical challenges.

##### In Vivo Studies

All animal procedures were approved by the Animal Use and Care Committee of Capital Medical University Affiliated Beijing Ditan Hospital and performed in accordance with institutional guidelines of Wuhan United Imaging Surgical Co., Ltd. (protocol No. ZR2025001; institutional policy 97400001-QER-RCA-01). Clinically healthy female Bama miniature pigs (∼30 kg) were used. Preoperative sedation was administered by intramuscular sufentanil (2.5 mg/kg) plus etomidate (1.5 mg/kg), and anesthesia was maintained with 2% isoflurane.

##### In Vivo Topical Hemostatic Tests

A 1-cm incision was made in the ear of an anesthetized piglet to induce active bleeding. Under white LED light (20 klux), the pre-gelled αLA–AdoB_12_ adhesive was applied to the bleeding site. Hemostasis was evaluated by monitoring bleeding cessation. The adhesive was subsequently removed using ethanol-soaked medical gauze.

##### In Vivo Endoscopic GI Hemostatic Tests

1. **Esophageal mucosal bleeding**. Superficial mucosal bleeding of an anesthetized piglet was created in the mid-esophagus using biopsy forceps (five pinches at the same site until capillary oozing was observed).
2. **Severe arterial intestinal bleeding**. Identified intestinal polyp of an anesthetized piglet was resected with a semicircular snare, resulting in brisk pulsatile bleeding.

For hemostasis, 1 mL of αLA–AdoB_12_ precursor solution was endoscopically delivered to the bleeding site via an endoscopic injection needle. Endoscopic white light illumination (∼20 sec) photostabilized the precursor to form an adhesive barrier. Hemostasis was evaluated by irrigating the wound with water through the injection needle to assess barrier stability under hydrodynamic stress.

### Supplementary Figures

**Fig. S1.**
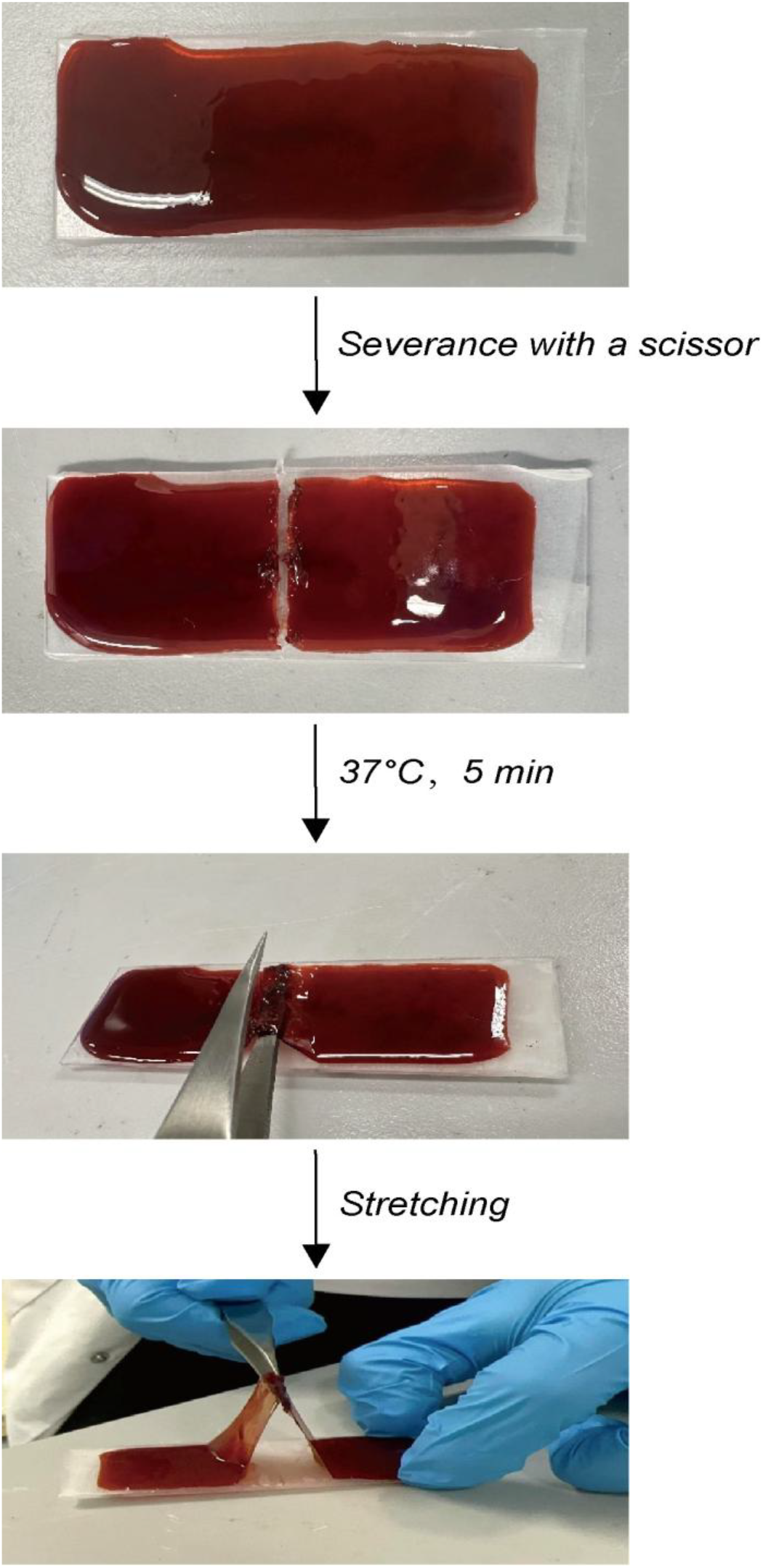
Self-healing of a photostabilized αLA-AdoB_12_ adhesive patch.

**Fig. S2.**
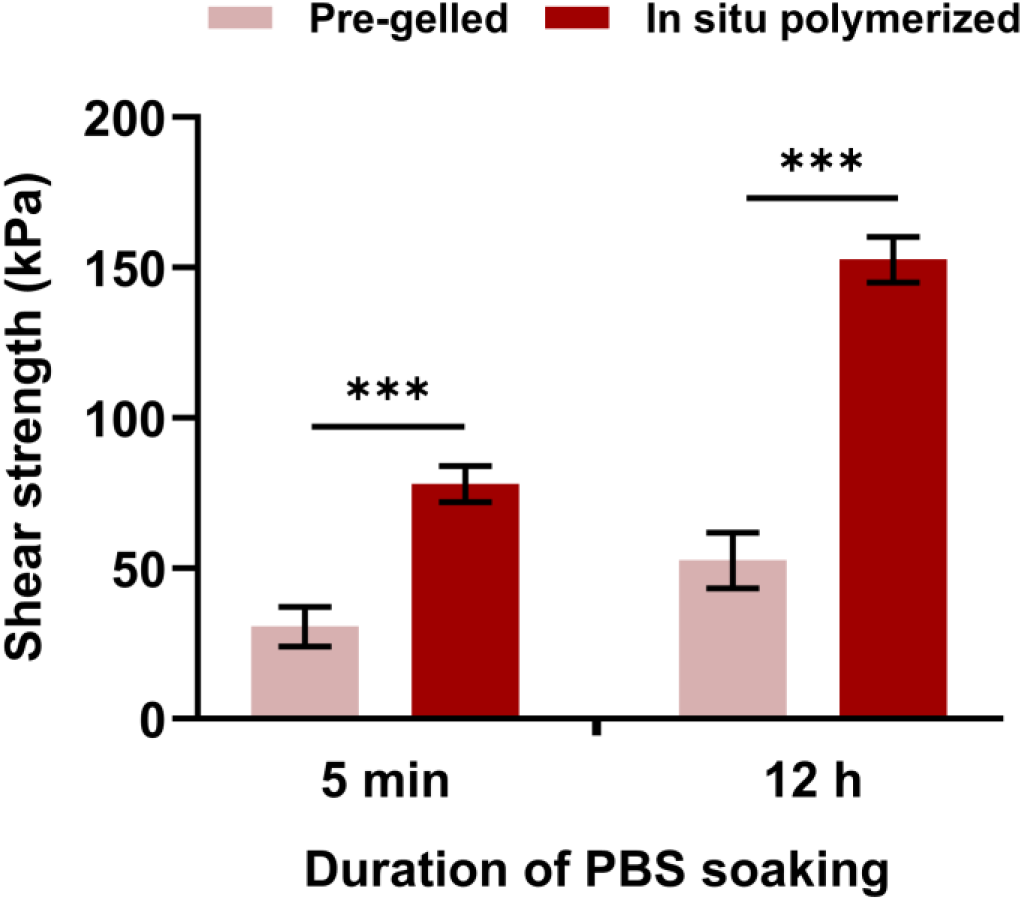
Ex vivo water-resistant adhesive performance of photostabilized poly(αLA)-AdoB_12_. Comparison of adhesion strength for pre-gelled patches vs. in situ polymerized αLA-AdoB_12_ adhesives after 5-min or 12-h PBS soaking. Statistical significance was determined by unpaired two-tailed t-test (***p ≤ 0.001).

**Fig. S3.**
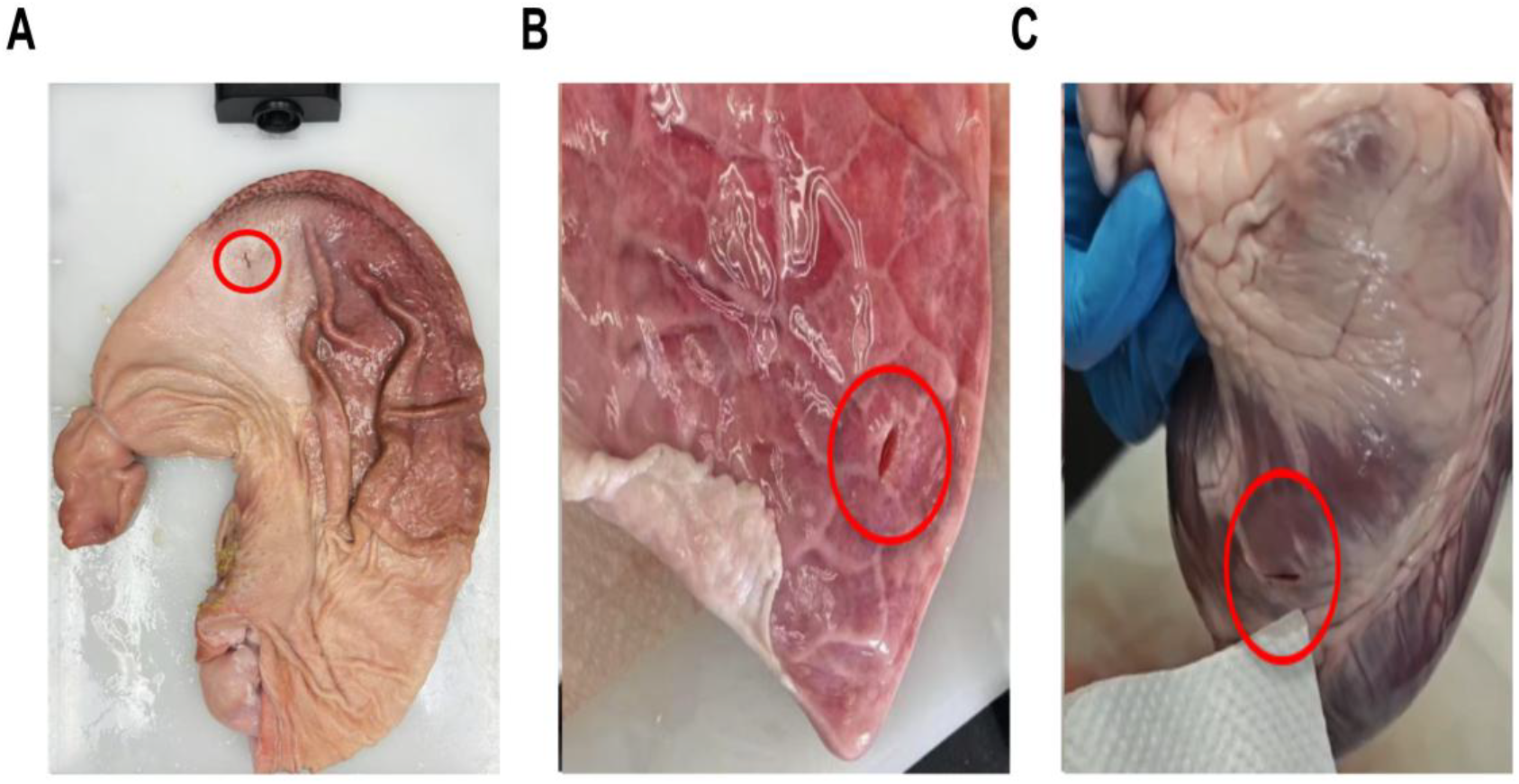
One-centimeter incisions made on porcine organs [stomach (A), lung (B), and heart (C)] using a scalpel.

**Fig. S4.**
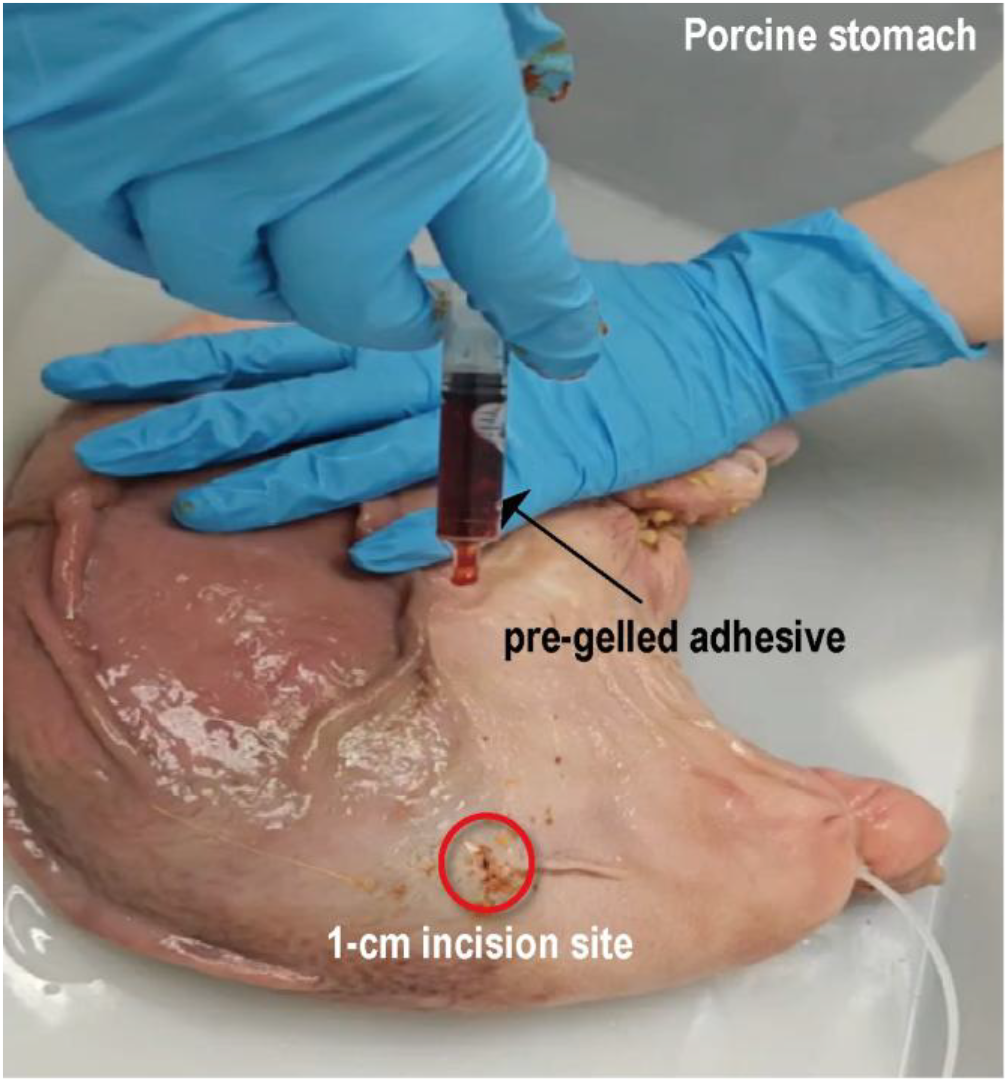
Application of pre-gelled αLA–AdoB_12_ adhesive. Precursor solution in syringe after 5-min white LED light irradiation (20 klux), achieving a syringe-deliverable viscosity for targeted deposition onto a 1-cm incision on porcine stomach.

**Fig. S5.**
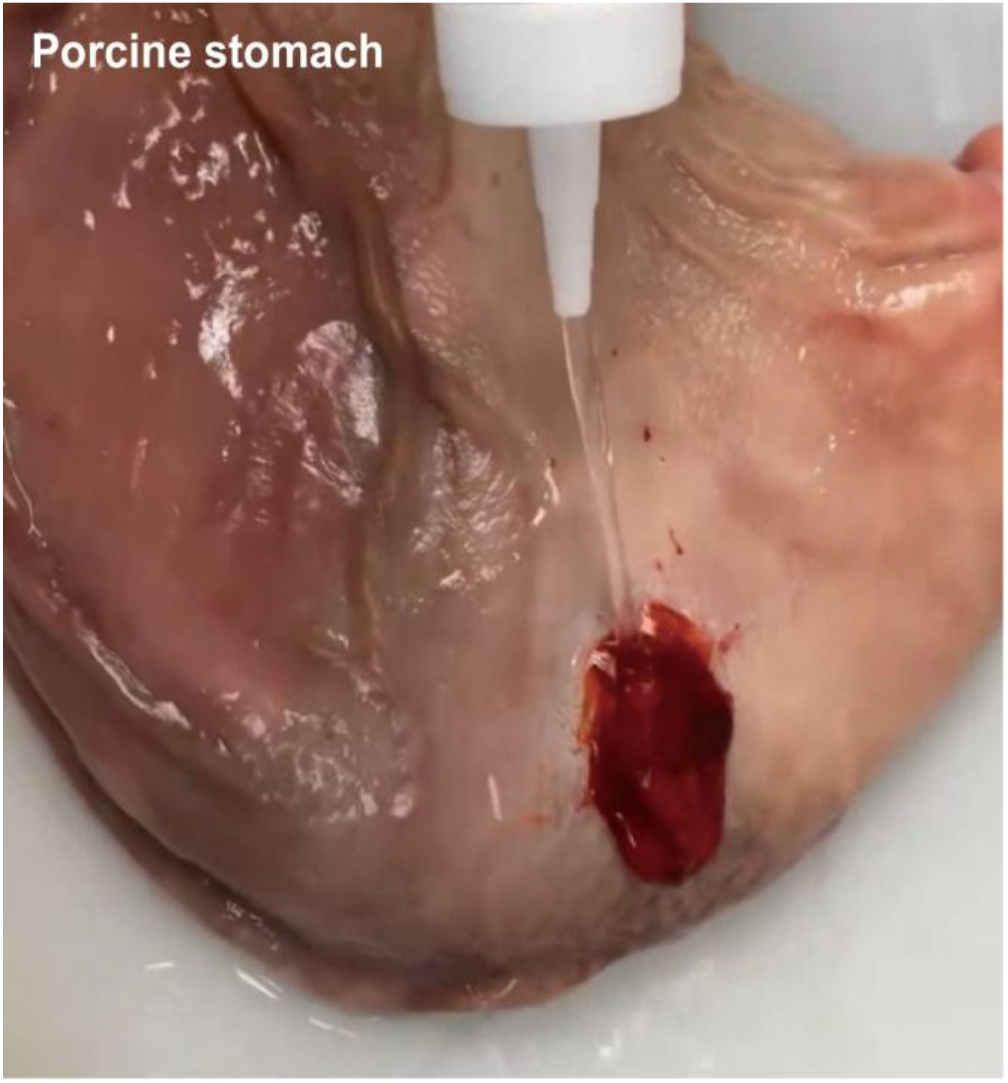
Hydrodynamic challenge of adhered pre-gelled αLA–AdoB12 adhesive on a porcine stomach. The adhesive seal remains intact under continuous water flushing.

**Fig. S6.**
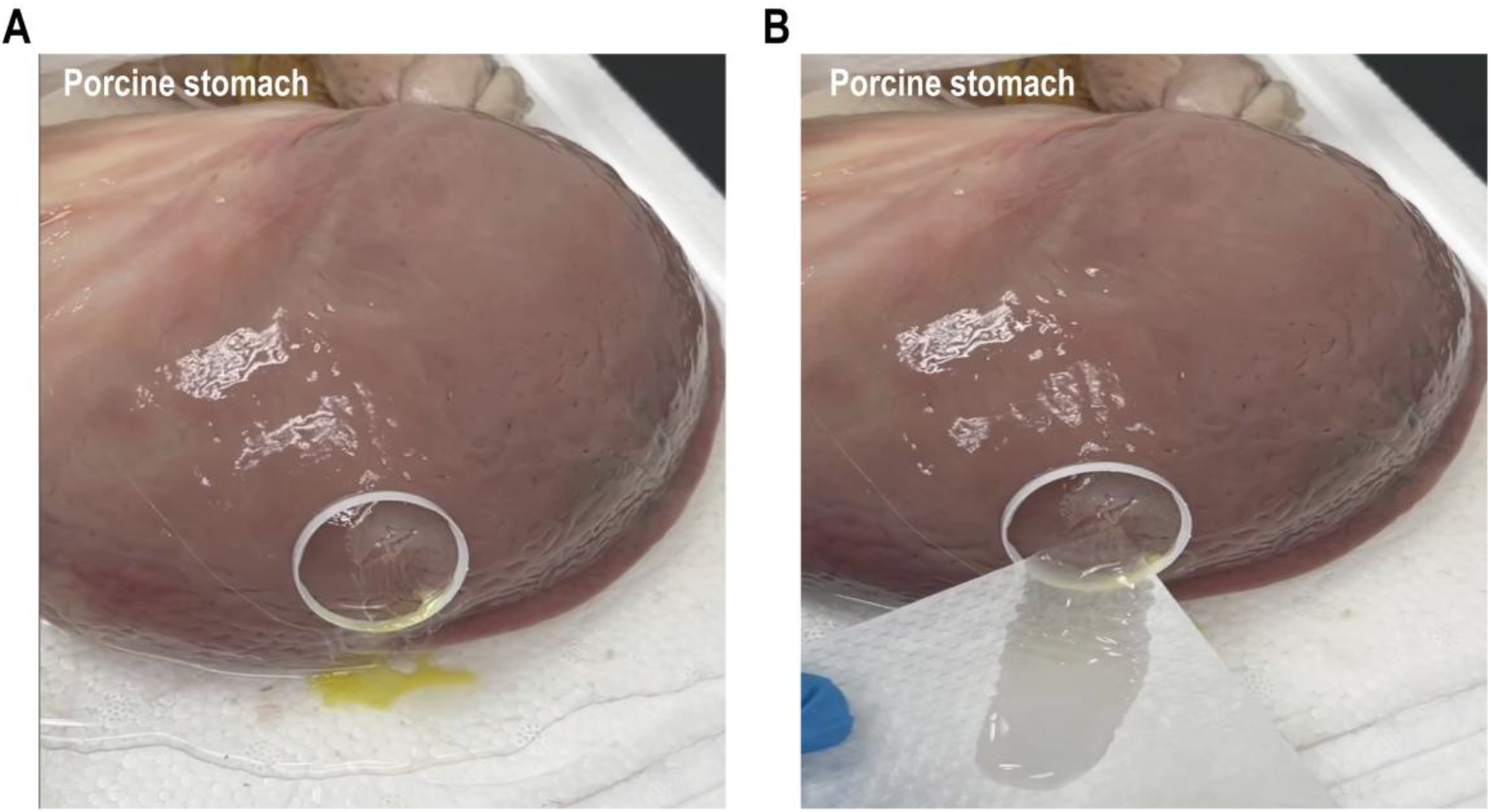
αLA alone fails to seal the incision site of a water-filled porcine stomach. **A**. The application of an αLA solution (yellow) to an incision site demonstrates its poor adhesive properties. **B**. Flowing water rapidly dispersed the solution, resulting in leakage. This control experiment underscores the necessity of a stabilizer, such as AdoB_12_, to achieve robust adhesion and effective sealing under wet, dynamic conditions.

#### Movie Descriptions

**Movie S1. Photo-induced gelation of αLA–AdoB**_**12**_ **under aerobic and anaerobic conditions**. Precursor solutions were irradiated with white LED light (20 klux, 5 min) in the presence or absence of oxygen. The video is shown at 5× real-time speed.

**Movie S2. Self-healing and stretchability of αLA–AdoB**_**12**_ **adhesive**. The video shows rapid self-healing, as well as full recovery of morphology within 5 minutes at 37°C.

**Movie S3. Ex vivo leakage from porcine lung without adhesive**. The video shows water leakage from a porcine lung incision in the absence of αLA–AdoB_12_, demonstrating baseline failure to retain fluid under perfusion.

**Movie S4. Ex vivo sealing of porcine lung with αLA–AdoB**_**12**_ **precursor**. The video depicts the sealing efficacy of the αLA–AdoB_12_ precursor solution under hydrodynamic conditions.

**Movie S5. Ex vivo leakage from porcine heart without adhesive**. The video illustrates water leakage from a porcine heart incision in the absence of αLA–AdoB_12_, establishing the unsealed control under flow.

**Movie S6. Ex vivo sealing of porcine heart with αLA–AdoB**_**12**_ **precursor**. The video demonstrates sealing of a porcine heart incision using the αLA–AdoB_12_ precursor solution, while highlighting challenges with precursor flowability in a dynamic, wet environment.

**Movie S7. Ex vivo leakage from porcine stomach without adhesive**. The video records water leakage from a porcine stomach incision in the absence of αLA–AdoB_12_, serving as the unsealed control.

**Movie S8. Ex vivo sealing of porcine stomach with pre-gelled αLA–AdoB**_**12**_ **(manual compression)**. The video demonstrates application of pre-gelled adhesive achieving a fluid-tight seal on a porcine stomach incision during manual pressing.

**Movie S9. Ex vivo sealing of porcine stomach with pre-gelled αLA–AdoB**_**12**_ **(continuous flushing)**. The video captures sustained fluid-tight sealing of a porcine stomach incision under continuous water flushing after application of pre-gelled adhesive.

**Movie S10. Failure of αLA alone in ex vivo stomach sealing**. The video reveals that αLA alone (without AdoB_12_) fails to prevent water leakage from a porcine stomach incision, underscoring the need for the αLA–AdoB_12_ system.

**Movie S11. Instant hemostasis triggered by blood contact**. The video captures rapid adhesion and conformal barrier formation upon delivery of the pre-gelled adhesive to a bleeding wound, resulting in immediate hemostasis.

**Movie S12. αLA–AdoB**_**12**_ **adhesive functions as a temporary hemostatic barrier**. The video showcases robust sealing, support of endogenous clot formation, and the possibility of gentle removal prior to clot maturation without disrupting the nascent clot.

**Movie S13. Endoscopic hemostasis of moderate esophageal bleeding in a porcine model**. The video demonstrates control of mucosal hemorrhage using light-activated adhesive, which withstands hemodynamic shear stress under irrigation.

**Movie S14. Stabilization of severe arterial bleeding via adhesive self-healing**. The video highlights the material’s resilience under high-flow bleeding, including self-healing behavior and enhanced stabilization after 20 seconds of endoscopic light exposure (20 klux).

